# A torsion-based rheometer for measuring viscoelastic material properties

**DOI:** 10.1101/2020.09.16.288415

**Authors:** Elise Jutzeler, Merrill Asp, Katherine Kerr, Dawei Song, Alison Patteson

## Abstract

Rheology and the study of viscoelastic materials is an integral part of engineering and the study of biophysical systems however the cost of a rheometer is only feasible for colleges, universities and research laboratories. Even if a rheometer can be purchased it is bulky and delicately calibrated limiting its usefulness to the laboratory itself. The design presented here is less than a tenth of the cost of a professional rheometer and portable making it the ideal solution for high school students as a way to introduce viscoelasticity at a younger age as well as for use in the field for obtaining preliminary rheological data.

## Introduction

K-12 science instruction is currently experiencing a revolution; a shift from a view of standards-based instruction as a checklist of items to be covered to a three-dimensional approach wherein content is covered while integrating instruction in cross-cutting concepts such as cause and effect and science and engineering practices like analyzing and interpreting data. As classroom teachers work to achieve this shift one problem remains; the existence of a disconnect between what high school students and college researchers do. The primary reason for this disconnect is the immense cost that surrounds the hi-tech equipment used in college and professional labs. The solution to this issue is the design and creation of an affordable and portable version of such materials that can be used by students to introduce them to advanced lab practices before they graduate from high school. Here, we present a low-cost portable rheometer, which can be used to measure the mechanical properties of many different materials, and present a biophysical application, in which the non-linear mechanical properties of tofu are investigated as a phantom for liver disease diagnosis.

## Background

There are many ways to categorize materials. One such way is as an elastic solid or a fluid. While these two types of materials are often covered in commencement level physics courses in the United States, they fail to address one important point; most materials exhibit a combination of these solid and fluid properties and are therefore characterized as viscoelastic. The study of such materials is an integral part of understanding structural stability in engineering, the importance of texture in food science, and the wear on joints in the body.

Whether a material is solid, or fluid, or some combination is often assessed by applying an external force on the material and evaluate its resulting deformation. Purely elastic solids behave as Hookean springs: the force *F* to stretch or compress the elastic solid is proportional to the amount of stretch of compression Δ*x*, *F* = *E*Δ*x*, where the elastic modulus of the material *E* acts like a spring constant. Fluids are frictional in nature. The force to deform a fluid is proportional to its velocity dx/dt, F = η d(Δ*x*)/dt, where η is the fluid viscosity.

Viscoelastic materials can be described in part like solids and in part like fluids in relation to how they move in response to applied forces (Larson,1999; Macosko, 1994). One such description is a Kelvin-Voigt material (Fig. 1), which models a viscoelastic material as a combination of an elastic spring and a movable plunger immersed in a viscous liquid (also known as a dashpot) as shown schematically in the diagram below.

**Fig. 1.**
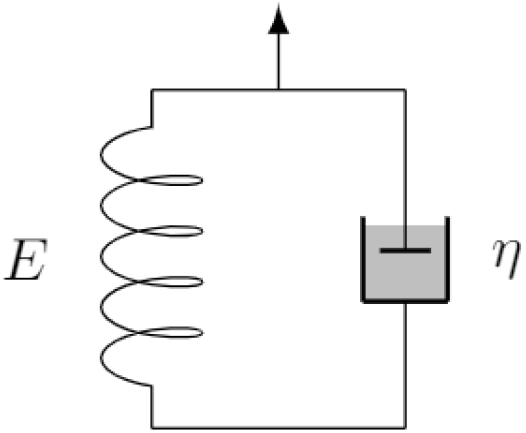
Kelvin-Voigt model of a viscoelastic material. The spring represents the elastic components of the material and the dashpot represents the viscous component.

The key feature of this model is that the two elements are combined in parallel. That is, the two elements have the same displacement Δ*x*, and the overall force on the material is the sum of the forces from each element, so F = *E*Δ*x* + η d(Δ*x*)/dt.

Oscillatory rheology is one of the most commonly used techniques to quantify the viscous and the elastic components of a material (Macosko, 1994) (Fig. 2). In this test, the material sample is driven by a plate undergoing an oscillatory displacement such that the graph of θ with time makes a sine wave, θ(t) = Asinωt, where ω is the oscillatory frequency and A is the wave amplitude. In a commercial rheometer, a force transducer measures the resulting stress from the sample as it is being deformed.

**Fig. 2.**
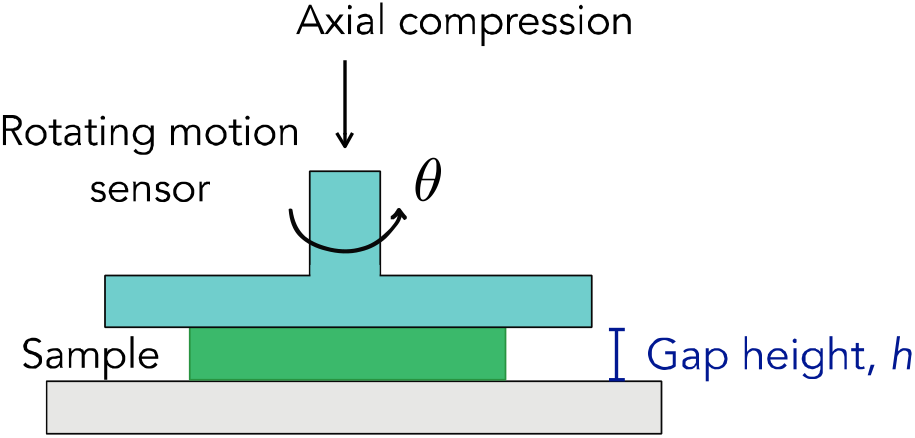
Schematic of a parallel-plate rheometer. The sample is wedged between two plates. The bottom plate is fixed, and the top plate rotates, allowing the application of shear stress. Axial compression or extension can be applied by changing the gap height.

For a solid material, the force is proportional to the displacement. Thus, the measured force as a function of time is completely in phase with the displacement of the material, which is oscillating at θ = Asinωt (Fig. 3). In contrast, the fluid material has a force that is proportional to rate of displacement dθ/dt = Aωcos(ωt). This wave is exactly 90° out of phase with the oscillations as the liquid is only under stress when the displacement is *changing.* As soon as the plate stops moving, the liquid dissipates any internal stress nearly instantly.

**Fig. 3.**
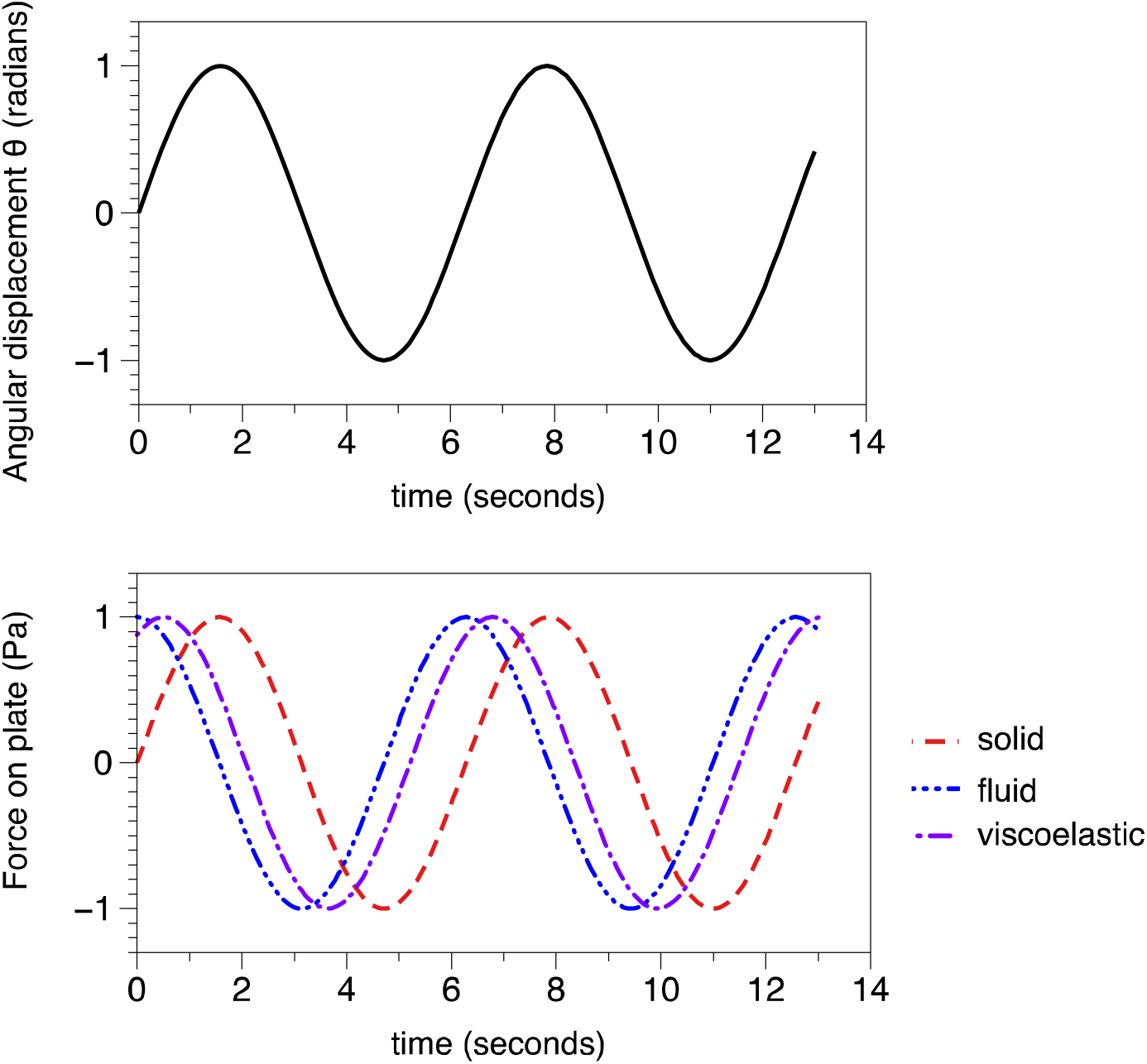
Force-displacement response of common materials under oscillatory shear.

Viscoelastic materials exert a force on the top plate that is in part proportional to the material displacement (sinωt) and the displacement rate (cos(ωt)), such that F = A sinωt + B cosωt, which can equivalently be written as F = C sin(ωt + δ), where δ is the phase angle or phase shift between the deformation and the material response. In solids, δ is very small and “in phase” with the oscillations. A perfectly elastic solid will have no delay at all, and stress will change simultaneously with strain. The quantity tan(δ) is a measure of how solid or fluid-like the material is. If tan(δ)≫1, the material is more fluid-like. If tan(δ)≪1, it is more solid-like.

In these oscillatory shear tests, we typically define the material properties in terms of stress and strains as opposed to forces and displacements. The stress σ is a measure of the forces (F/A) exerted on the plate by the material when it is deformed. The strain γ is a measure of the amount of deformation a material undergoes as a result of an applied force. In an oscillatory test, the strain is proportional to the angular position θ of the rheometer plate at any given instant and the dimensions of the sample. We can then define the storage modulus, G’, which characterizes the solid-like component of the material and the loss modulus, and G’’, which characterizes the fluid-like component, such that the stress is given by

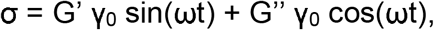

and tan(δ) = G”/G’. The “storage” and “loss” refer to the storage and loss of *energy* in the material. Energy can be stored in an elastic material by compressing or stretching it, whereas mechanical energy is dissipated into heat – essentially lost – when a plunger moves through liquid.

Commercial rheometers rely on expensive force transducers to determine the torque associated with sample strain and can cost upward of $40,000-$200,000. Here, we present a portal version that can be assembled for less than $400, making it much more practical for field use and educational purposes.

## Materials and Methods

Our torsion-based rheometer stems from a slightly altered version of the oscillatory test and is related to the torsion pendulum developed previously (Janmey, 1991; Plazek 1958). Here, the top plate is comprised of a Vernier rotational motion sensor (Fig. 4). As opposed to applying a driven oscillation of the plate and trying to measure the resulting force, we simply apply a small perturbation to the plate and measure the resulting motion θ of the plate. In this test, the elastic solid component of the material acts to oscillate the sample back and forth while the fluid viscosity on the sample acts to dampen its motion. A typical θ versus time curve is shown in Fig. 5. This motion is described by a slightly-damped harmonic oscillator, owing to the spring and dashpot components that represent the materials response (Kelvin-Voigt model). The parameters G’ and G’’ can be estimated from the frequency (ω) and amplitude decay (Δ) of the θ versus time curve, as follows:

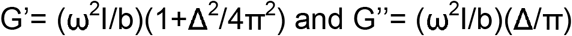

where I is the moment of inertia of the rotating plate and b is determined by the radius R and height h of the test sample by the formula b = πR^4^/2h. A description of these is available in (Staverman, et al, 1956) for German readers. For convenience, we provide a full derivation of these equations in the Supplementary Materials, which should be appropriate for undergraduate Physics students in a Vibrations and Waves course, for instance.

**Fig. 4.**
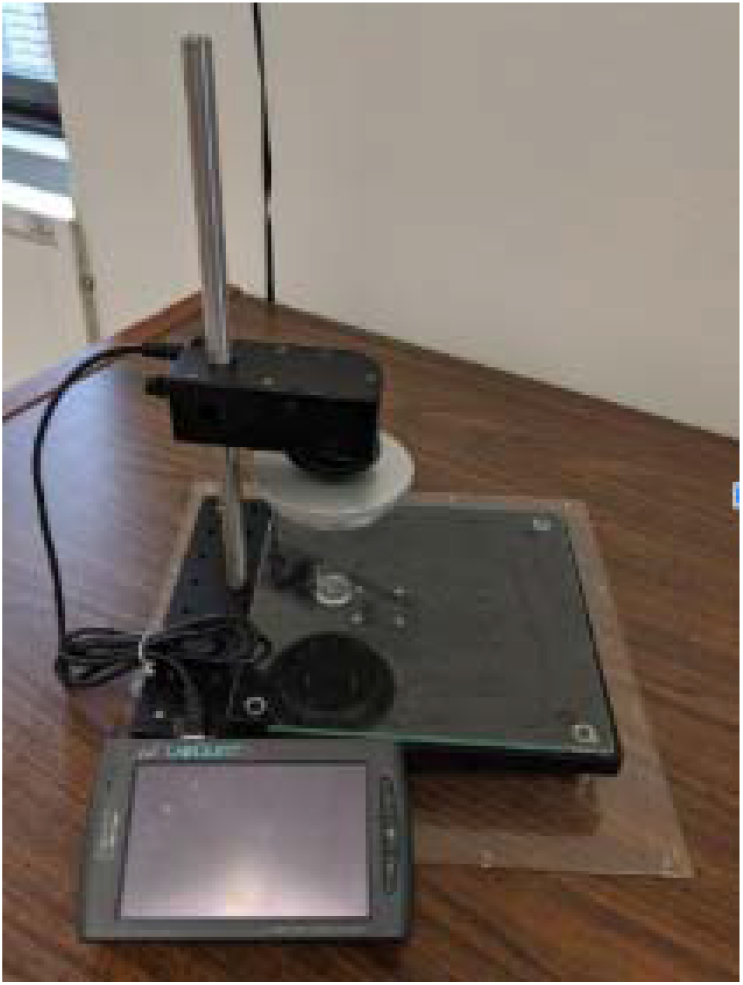
Mounted Vernier sensor with PDMS sample.

**Fig. 5.**
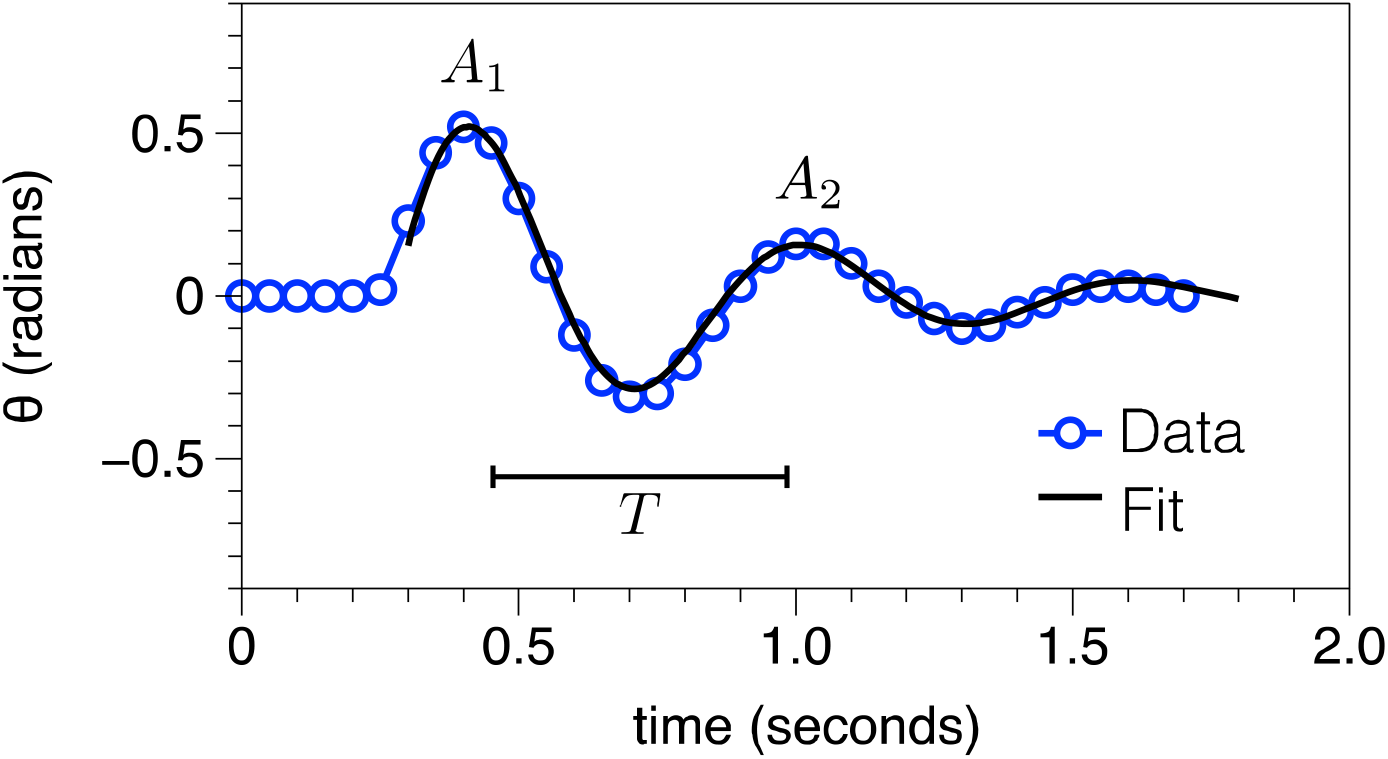
JELL-O Jiggler output showing angular displacement as a function of time. The blue circles are data points from the Rotary Motion Sensor and the black line is the fit to a damped harmonic oscillator *θ*(*t*) = *A*_1_ cos(*ωt)* exp (−*t*/τ).

The Vernier Rotary Motion Sensor (RMV-BTD) and Accessory Kit (AK-RMV) (Fig. 4) used in this design consists of a number of aluminum and steel disks allowing students to alter the moment of inertia of the rotating plate. The kit also comes with a pulley attachment allowing for other rotation related experiments. The sensor is attached to a vertical pole mounted on an optical plate (Fig. 4) and then lowered to sandwich the test sample between the sensor’s rotating plate and a sheet of plate glass.

As described the apparatus cost $379.76.

| Manufacturer | Item ID | Description | Cost |
| --- | --- | --- | --- |
| Thorlabs | MB8 | Solid Aluminum Optical Breadboard 8x8x1/2 | \$77.22 |
| | DF1/M | Delrin Breadboard Feet | \$21.54 |
| Vernier | Rotary Motion Sensor | RMV-BTD | \$169 |
| | Rotation Motion Accessory Kit | AK-RMV | \$112 |
| | LabQuest 2 (Optional) | LABQ2 | \$339 |

This design works best with soft samples such as tofu or JELL-O making it the ideal equipment for introducing viscoelastic properties as they relate to the field of food science or biomaterials. Materials which are too firm will slip or skid during testing. Ideal test samples should be cut into a circular disk no larger in diameter than the plate itself. Once the sample is in place a small torque is manually applied to the rotating plate and the oscillations of the plate are recorded on a Lab Quest 2 or other compatible devices.

Without a test sample the plate would simply rotate in one direction eventually coming to rest due to friction within the sensor’s hardware however with the addition of the sample the resulting motion is that of a damped oscillator (Fig 5). Analysis of the oscillation frequency and the logarithmic decrement allow for a calculation of the sample’s storage and loss moduli.

## Results and Discussion

This classroom rheometer was tested on three soft materials; PDMS, Firm Tofu, and JELL-O Jigglers. Jigglers are a stiffer gelatin recipe with a higher gelatin to water ratio.

**Table 1.** Table showing results from the classroom rheometer design as well as the laboratory rheometer.

| Material | Specifications | Classroom $G'$ (kPa) | Malvern Results (kPa) | Classroom $G''$ (kPa) | Malvern Results (kPa) |
| --- | --- | --- | --- | --- | --- |
| PDMS | SYLGARD 527 1:1 | 0.83 | 1.11 | 0.34 | 0.54 |
| Tofu | Firm | 23 | 20. | 8.2 | 6.4 |
| JELL-O Jigglers | According to Pkg. | 1.6 | 1.1 | 0.19 | 0.21 |

The motivation for this design was to create an affordable way to bring rheometry into the classroom and at less than $400 that goal can be a reality. Additional modifications can be made to decrease costs even more. A standard classroom ring stand and piece of glass from a picture frame or old window could easily be substituted as a base to hold the apparatus and the Lab Quest 2 can be replaced by classroom computers or other compatible devices. Furthermore, the implementation of the design is flexible allowing teachers options as to how and when they introduce it to their classrooms. Students can use the sensor to determine the moduli for a particular material such as JELL-O Jigglers as an isolated lab experiment or they can use the equipment throughout the course of the school year as part of a larger research assignment. While studying rotational motion they can calculate the moment of inertia for the different plates and then experimentally determine them using the sensor. They can also experiment with different JELL-O recipes to determine the stiffness that returns the best oscillation data. Then during the unit on waves, they can interpret displacement vs. time data to determine the frequency and logarithmic decrement. A sample NGSS aligned lesson can be found in the supplemental materials of this paper.

### Biophysics Application: Viscoelastic properties of tofu as a phantom for liver disease

Emerging studies are increasingly highlighting the importance of mechanical forces on biophysical systems (Thompson, 1942; Discher, et al 2005). For instance, the mechanical properties of tissues regulate many cellular functions, including motility, differentiation, and proliferation (Janmey, et al. 2011). Proper maintenance of tissue rigidity is thus a crucial aspect of healthy tissue function, and dysregulation of tissue rigidity is associated with many diseases, such as fibrosis and some forms of cancer (Bonnans, et al. 2014).

Liver disease is a salient example of a disease associated with changes in the mechanical properties of the tissue and is the cause of two million deaths annually (Asrani, et al. 2019). In this case, tissue stiffness is used as a diagnosis of liver disease progression (Mueller, et al. 2010). Tissue stiffness is typically assessed in patients using noninvasive techniques, such as magnetic resonance elastography (MRE) that uses mechanical waves to estimate the shear modulus of the liver tissue (Chundru, et. al. 2014).

The shear modulus is just one metric to assess tissue mechanical properties. Real tissues are viscoelastic solids. Their properties are also a function of the applied deformation. For instance, liver tissue softens with increasing applied strain (See comment *). In addition to shear strain, growing work is documenting the response of tissue to compressive loading. A recent report (Perepelyuk et, al. 2016) showed that rat liver tissue stiffens upon uniaxial compression (Fig. 6a), and that this compression-stiffening behavior was significantly stronger in fibrotic liver tissue compared to normal liver tissue. These results highlight how diseases can alter how tissues respond to forces.

**Fig. 6.**
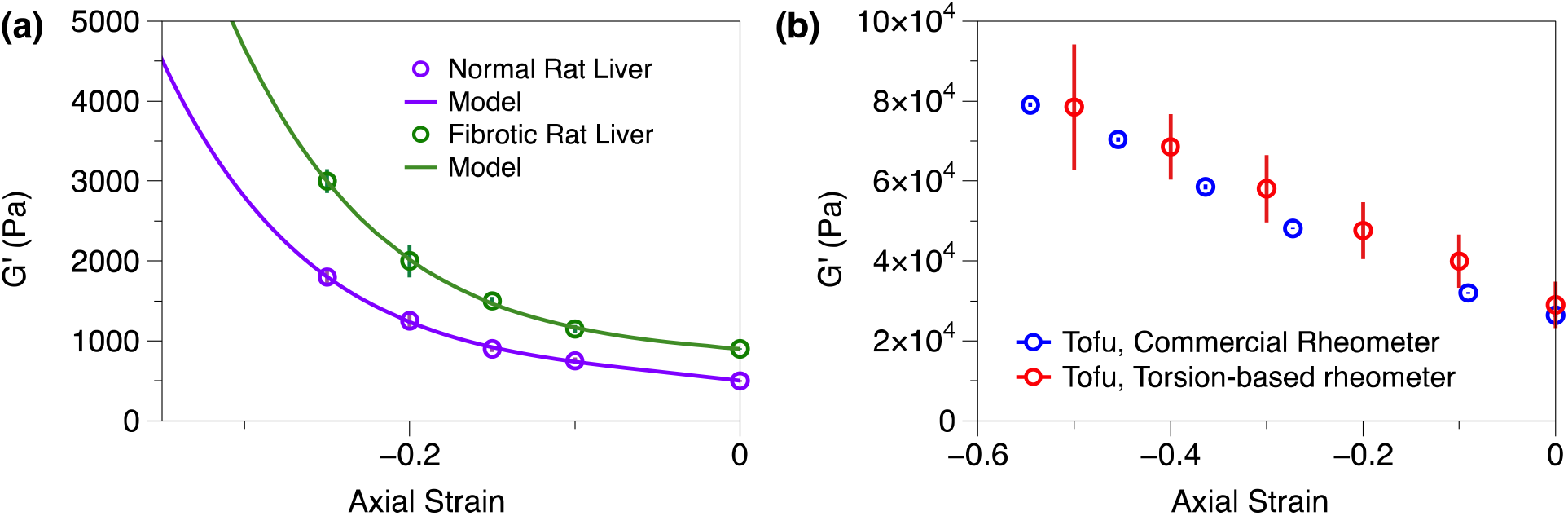
Compression stiffening behavior of liver tissue and tofu samples. (a) The shear modulus G’ increases upon axial compression in both normal and fibrotic rat livers, adapted from Perepelyuk et, al. The solid lines are fits to the data using the Ogden hyperelastic model. (b) Tofu exhibits the compression stiffening behavior observed in liver samples. Data shows comparison between data obtained in torsion-based rheometer ($400) vs. a commercial Kinexus rheometer ($100k+).

The highly non-linear properties of liver tissue are attributed to complex material features, such as the ability of water to flow through the tissue and the ability of cells to form dynamic connections with the extracellular polymeric matrix of the tissue. While the requirements of high school coursework may be limited to uniaxial linear systems the introduction of this complex case even if only conceptually has the potential to provide context for student learner and foster interest in future study of these and other biomedical applications.

Experiments with real tissues are impracticable in high school and undergraduate settings, but there exist tissue simulating phantoms worth studying in their own right. Tissue phantoms are of practical interest, allowing for the development and calibration of MRE devices, for instance. Here we demonstrate that one of the most easily accessible tissue phantoms, tofu (Belmont, et al 2013; Wu, 2001), mimics the compression-stiffening behavior of real liver tissue and that our custom bench-top rheometer can be used to accurately capture the phenomena.

To perform a compression test, we substitute our fixed bottom plate for an adjustable stand. To begin, the tofu sample is placed on the stand, and the stand is raised just to the point that the tofu sample just makes contact with the top plate. This setting corresponds to a gap height that matches the initial height of the sample and a 0% compressional strain. The value of G’ and G” is then determined as described in Methods. Next, the tofu is compressed in a series of steps with G’ and G” being measured along each step of the way. Here, we compress the sample in 10% intervals by manually raising the position of the lower plate with the stand.

Figure 6b shows the results of Trader Joe’s Super Firm tofu in our custom rheometer device. We find that similar to liver tissue the shear modulus G’ of tofu increases upon compression. Also shown in Figure 6b are the G’ data of tofu gathered by a Kinexus Malvern commercial rheometer. These data were gathered from oscillatory tests using 2% shear strain and 1 Hertz frequency and computer software that allows automation of the gap height. Given the rather simple and crude design of our torsion-based rheometer, the torsion-based rheometers results are in remarkable agreement with those from the commercial rheometer.

In addition to experimental characterization, developing reliable models for tissue mechanical properties is of crucial importance, since it can provide valuable insights into the mechanisms involved in tissue deformation, and predict the response and evolution of living tissues in physiological and pathological conditions (Fung, 2013). Continuum mechanics-based models (Gurtin, et al. 2010) are especially suitable for describing the macroscopic response of tissue samples used in rheological measurements. Among those, Ogden hyperelasticity model (Ogden, 1972) has been shown to be successful in describing the rheology of soft tissues like liver, brain and fat (e.g., Gao, et al. 2010; Mihai, et al. 2015). In particular, it is able to accurately capture the highly nonlinear, compression stiffening behavior of liver tissue (Fig. 6a), after appropriately calibrating model parameters using available experimental data following the procedure of Mihai, et al. (2015). While a thorough understanding of Ogden model would require a strong background in continuum mechanics that is beyond the scope of high-school coursework, a brief introduction of these modeling concepts could stimulate students’ interests in pursuing further studies in the field of biomechanics and biophysics.

## Conclusion

This article presents a cost effective and portable way to assess the mechanical properties of materials. This tool can be used to incorporate viscoelastic materials and bio-physics concepts into high school level classes. The skills obtained in these laboratory activities provide students with a foundation that can better prepare students for their college coursework and future lab experiences.

## Supplemental Materials

See attached.

## Acknowledgements

We acknowledge useful conversations with Sam Sampere and Paul Janmey. This work was funded by NSF MCB 2026747.

## Author contributions

E.J. designed the torsion-based rheometer, performed and analyzed experiments, and wrote the lesson plan. M.A. derived analytical results. K. K. and D.S. designed and analyzed experiments with tofu. D.S. developed models to characterize liver mechanical properties. A.P. planned and analyzed experiments. E.J. and A.P wrote the paper.

* This behavior deviates from the model of a linear Hookean spring. Instead, it is suggestive of a model in which the spring constant is a decreasing function of strain.

## Notes

### Competing Interest Statement

The authors have declared no competing interest.

